# Evidence for a Recombinant Origin of HIV-1 group M from Genomic Variation

**DOI:** 10.1101/364075

**Authors:** Abayomi S Olabode, Mariano Avino, Tammy Ng, Faisal Abu-Sardanah, David W Dick, Art FY Poon

## Abstract

Reconstructing the early dynamics of the HIV-1 pandemic can provide crucial insights into the socioeconomic drivers of emerging infectious diseases in human populations, including the roles of urbanization and transportation networks. Current evidence indicates that the global pandemic comprising almost entirely of HIV-1/M originated around the 1920s in central Africa. However, these estimates are based on molecular clock estimates that are assumed to apply uniformly across the virus genome. There is growing evidence that recombination has played a significant role in the early history of the HIV-1 pandemic, such that different regions of the HIV-1 genome have different evolutionary histories. In this study, we have conducted a dated-tip analysis of all near full-length HIV-1/M genome sequences that were published in the GenBank database. We used a sliding window approach similar to the ‘bootscanning’ method for detecting breakpoints in intersubtype recombinant sequences. We found evidence of substantial variation in estimated root dates among windows, with an estimated mean time to the most recent common ancestor (tMRCA) of 1922. Estimates were significantly autocorrelated, which was more consistent with an early recombination event than with stochastic error variation in phylogenetic reconstruction and dating analyses. A piecewise regression analysis supported the existence of at least one recombination breakpoint in the HIV-1/M genome with interval-specific means around 1929 and 1913, respectively. This analysis demonstrates that a sliding window approach can accommodate early recombination events outside the established nomenclature of HIV-1/M subtypes, although it is difficult to incorporate the earliest available samples due to their limited genome coverage.

## Introduction

At present, nearly 37 million individuals are living with human immunodeficiency virus type 1 (HIV-1) worldwide [1]. Reconstructing the early dynamics of the HIV-1 pandemic can provide crucial insights into the socioeconomic drivers of emerging infectious diseases in human populations, including the roles of urbanization and transportation networks [2]. Current evidence indicates that the global pandemic comprising almost entirely of HIV-1/M originated around the 1920s in central Africa [2], where it had already diversified into the major subtypes before spreading around the world to their present-day distribution [3, 4]. Much of the support for this model on the origin of HIV-1/M is based on the analysis of contemporary sequence diversity in specific regions of the HIV-1 genome [5]. Due to the rapid evolution of this retrovirus, genetic differences accumulate on a directly observable time scale, making it possible to estimate the rate of molecular evolution, *e.g*., molecular clock, from sequences that were sampled at different points in time [6], even when they are separated by only a few decades. In practice, these ‘dated-tip’ analyses of HIV-1/M diversity often rely on a number of assumptions including an absence of recombination, such that the entire span of the multiple sequence alignment can be related through a single phylogeny [2, 3, 5]. Previous studies have shown that recombination can induce skew estimates of the molecular clock [7] and thereby estimates of times to common ancestors. For example, the transfer of divergent sequence by recombination from another subtype could inflate molecular clock estimates within that interval.

HIV-1/M is subdivided into nine major subtypes [8] and at least 90 circulating recombinant forms (CRFs) [9], which are defined as recombinants between the recognized subtypes that have been observed in three or more persons who are not epidemiologically linked [10]. Some CRFs have become highly prevalent in specific regions of the world, such as CRF01_AE in southeast Asia and CRF02_AG in west Africa [4]. Homologous recombination from template-switching events during reverse-transcription is an obligate step in the replication cycle of HIV-1 [11] and estimates of the recombination rate *in vivo* are on the same order of magnitude as the rate of mutation [12]. Thus, new recombinant forms are constantly emerging and inter-subtype recombination has recently been estimated to have occurred in approximately 20% of HIV-1 lineages over the last 30 years of the global pandemic [13]. There is growing evidence that recombination has played a significant role in the early history of the HIV-1 pandemic. For instance, several groups have described novel recombinant forms that incorporate genomic fragments from highly divergent lineages that precede the proliferation of the currently recognized HIV-1/M subtypes [14–16]. These observations suggest that what we presently recognize as ‘pure’ subtypes are quite likely recombinants of old, rare or extinct lineages, as illustrated by CRF01_AE and its extinct parental lineage that was originally designated subtype E [17].

In this study, we have conducted a dated-tip analysis of all near full-length HIV-1/M genome sequences published in the GenBank database. We used a sliding window approach similar to the ‘bootscanning’ method for detecting breakpoints in intersubtype recombinant sequences [18], where a series of phylogenies are reconstructed on short intervals of the multiple sequence alignment. Instead of finding breakpoints, however, we use dated-tip methods to locate the root in each phylogeny and then use recently developed maximum-likelihood methods [19] to rescale the tree in chronological time. The intended purpose of this ‘rootscanning’ approach is to mitigate the effect of inter-subtype recombination on estimates of the molecular clock and the time to the most recent common ancestor. We describe a network-based strategy to overcome the inherent difficulties in generating a multiple alignment of divergent HIV-1 genome sequences that include extensive insertion-deletion (indel) variation. Our analysis is consistent with a recombinant origin of HIV-1/M diversity and supports a reassessment of the current HIV subtype nomenclature in the era of next-generation sequencing where full-length genomes are rapidly accumulating on a global scale [20].

## Methods

### Data Collection

We queried the GenBank nucleotide database using the search term “Human immunodeficiency virus 1”[porgn:_txid11676] and limited the results to records with a minimum sequence length of 8000 nt. This query yielded a total of 7,816 records on October 11, 2017. We manually screened these records for laboratory clones, *in vitro* recombinants, non-group M variants and multiple samples from the same individual, in which case we retained only the earliest sample. For any sample with a missing collection date, we manually evaluated the publications associated with the GenBank record and restored the date information whenever possible; otherwise the record was discarded. After these steps, the resulting database comprised 3,900 records.

### Sequence Alignment and Processing

Phylogenetic reconstruction and extrapolation of times to the most recent common ancestor requires the alignment of homologous nucleotides in the respective sequences. Using a standard multiple sequence alignment program was not feasible for our purpose because the resulting alignment would contain an excessive number of gaps (see Supplementary Material, Fig. S1), due to the extensive numbers of substitutions and indel events among sequences in HIV-1/M, *i.e*., between defined subtypes and circulating recombinant forms. Hence, we needed to filter the genome sequences for nucleotide sites retaining a relatively high degree of evolutionary homology. We adopted a pairwise alignment approach against a single reference genome sequence in which we systematically excised problematic sites. By removing any insertions relative to this modified reference, the resulting pairwise alignment will strictly retain only sites that are homologous to the reference.

First we needed to generate an appropriate reference genome to carry out this Procrustean approach to sequence alignment. We generated a consensus sequence from a multiple alignment of a subset of genomes selected to be representative of the overall diversity in HIV-1/M. For this purpose, we used a p-spectrum kernel [21] to generate a pairwise distance matrix from the entire set of unaligned genome sequences. Specifically, we used a Python script to count the frequencies of all possible hexamers (*p* = 6) in each sequence and then calculated the inner product for every pair of frequency vectors:

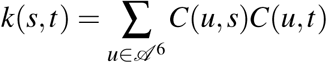

where 𝒜 = {*A*, *C*, *G*, *T*} and *C*(*x*, *y*) is a function that counts the number of instances of substring *x* in string *y*. We used the standard normalization 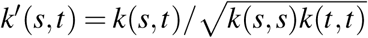 such that 0 ≤ *k*′ ≤ 1 [21]. The resulting similarity matrix was then converted into an undirected adjacency graph using a threshold of 0.93. We used the community detection algorithm called the *cluster* _*Jeading*_*eigen()* in the *igraph R* package to identify communities within the graph. For communities comprising of greater than 4 members, one member having the highest degree centrality (connectivity) in each community was selected, giving rise to 32 sequences. Having a higher degree simply means that the sequence is highly connected with other sequences when compared with others [22]. The selected 32 representative sequences were aligned using MAFTT 9 version 7.271) [23] to generate a multiple sequence alignment (MSA). We then generated a plurality-rule consensus sequence of 8921 nt from the MSA. Next, we used an implementation of the Altschul-Erickson algorithm as a C extension for Python (https://github.com/ArtPoon/gotoh2) to generate pairwise alignments of all sequences against this consensus sequence, discarding any insertions relative to the consensus to yield a draft alignment. Finally, we generated a MSA of 10171 nt from the draft alignment using MAFFT, then manually filtered residual regions with low homology (frequent indels) by removing alignment columns with a gap frequency greater than 70%. This led to the removal of 1544 columns to get a final alignment of 8627 nt. To evaluate the impact of the number of genomes on run time and the length of the resulting alignment, we drew random samples of sequences without replacement to build sub-alignments ranging from 100 to 3,500 sequences at increments of 200 sequences, using MAFFT with 12 threads.

### Phylogenetics and Molecular Clock Analysis

Windows of 500 nt were extracted at steps of 100 nt from the final alignment leading to generation of 82 unique windows. Each window was visually inspected for residual alignment errors and problematic sites, which were manually removed. For each window, we reconstructed a maximum likelihood (ML) phylogenetic tree using FastTree2 [24] under the default general time-reversible (GTR) model. Each tree was rooted based on the sample collection dates and a strict molecular clock model using the root-to-tip function (*rtt*) in the *ape* package in *R*. We used least-squares dating (LSD) [19] to rescale each rooted ML tree in time under a relaxed clock model, from which we estimated the time to the most recent common ancestor (tMRCA) for each tree. In cases where the sample collection dates were reported at the reduced precision of years or months, we input the date for the corresponding tip as a range; for example, the entry ‘2009-06’ was input as ‘b(2009.414,2009.493)’. To calculate the confidence intervals associated with the estimated tM-RCA and rate, the LSD program uses a parametric bootstrap approach where the branch lengths in the tree are resampled from a Poisson distribution calibrated to the number of sites. We set the sampling number to 100 for this step. To characterize the autocorrelation in estimates of tMRCA across windows, we fit a piecewise linear regression model with two segments and zero slopes, and selected the best segment breakpoint using the Akaike information criterion. We measured for autocorrelation in the residuals of the regression analysis using the Durbin-Watson test as implemented in the *car* package in *R*.

To corroborate our results from least-squares dating, we analyzed the same data using a Bayesian sampling method implemented in BEAST (version 1.8.4) [25]. Since it is not feasible to generate a random sample from the posterior distribution for trees relating to thousands of sequences, we uniformly down-sampled the sequence alignments in each window with respect to time, by selecting up to 10 sequences per collection year at random without replacement. A previous simulation study [26] determined that uniform sampling of sequences over time can provide unbiased reconstructions of past dynamics. This down-sampling resulted in a median of 294 sequences (range 284-294), which in our experience is at the upper limit of sample size at which a chain sample in BEAST might be expected to converge to the posterior distribution in a reasonable amount of time. We used a Python script to generate BEAST XML files from a model template for all 82 windows and ran the analyses in parallel on a local computing cluster. Based on previous work [27], the model template was configured to use the Tamura-Nei (TN93) nucleotide substitution model with rate variation approximated by a discretized gamma distribution with 4 rate categories, an uncorrelated lognormal molecular clock and a ‘skyline’ coalescent model [28] with the default 10 population size categories. Each chain sample was propagated for 10^8^ steps; the first 10^7^ steps were discarded as burn-in and the remainder was thinned at intervals of 10^4^ steps.

### Selection Analysis

We used the Fast Unconstrained Bayesian Approximation (FUBAR) method as implemented in HyPhy version 2.3.1 [29] to identify codon sites undergoing either purifying or diversifying selection over the evolutionary history of HIV-1/M. FUBAR fits a limited number of rate categories defining non-synonymous (dN) and synonymous (dS) substitution rate combinations to individual codon sites for a given codon alignment and phylogenetic tree. We extracted codon alignments from the regions encoding *gag*, *pol* and *env* in our final sequence alignment obtained as described in the preceding section (Data Processing). Because the reading frames of these major genes were disrupted by our filtering of extensive indel polymorphisms from the final alignment, we limited the codon alignments to cover a portion of each gene comparable to a single window in our dated-tip analysis. The resulting codon alignment each comprised 396 nt (132 codons) with the following consensus sequence coordinates: *gag* 793 - 1197, *pol* 2085 - 2486 and *env* 6225 - 6623. Next, we used FastTree2 to reconstruct phylogenetic trees for each codon alignments.

## Results

We retrieved 7,816 near full-length HIV-1 genome records from Genbank. After discarding irrelevant, incomplete or redundant sequences and restoring missing sample collection dates from publications associated with the respective records, we were left with 3,900 HIV-1/M genome sequences. Based on results from the SCUEAL subtyping program [30], 51% of the sequences were classified as “pure” subtypes while 49% were classified as recombinant forms. The sequences were obtained from 86 different countries with a mean of 45 (range 1 – 460) sequences per country. The countries with the highest number of near full-length genome sequence records included United States (*n* = 1752), Thailand (*n* = 719), South Africa (*n* = 698), China (*n* = 490) and Brazil (*n* = 414). The mean length of the genome sequences was 8881 nt (range 8000 – 9897 nt). Sample collection dates associated with the sequences ranged between the years 1978 to 2016, with a mean of 108 (range 1 – 443) sequences per year.

Although it was feasible to construct a multiple sequence alignment from these data using a conventional program (*e.g*., MAFFT) in a reasonable amount of time (Supplementary Fig. S2), the resulting alignment became long and sparse due to an excessive number of gaps induced by highly divergent sequence intervals or non-homologous sequence insertions (Supplementary Fig. S3, top). Supplementary Fig. S1 summarizes the increasing alignment length with progressively larger numbers of genome sequences in the alignment, starting from a mean of 10,978 nt with 100 randomly selected sequences to a mean of 24,146 nt with 2900 sequences. As a result, we constructed a consensus genome sequence from the multiple alignment of a subset of representative genomes, which we selected by a network clustering analysis of a p-spectrum kernel matrix [21] of the entire data set. In brief, we calculated the inner products of hexamer frequencies for all pairs of genomes and converted the resulting matrix into a graph using a cutoff value to define edges. We used a clustering method to extract 32 subgraphs and identified the highest degree node in each subgraph. Fig. 1 illustrates the entire graph with the identified representative nodes emerging from the 32 largest communities. Finally, we screened the multiple sequence alignment of genomes corresponding to these central nodes for regions of low homology and generated the consensus from the remaining sites. Compared to the HXB2 reference sequences the *gag, pol and env* genes of the consensus sequence are 1475 (HXB2 1503), 3000 (HXB2 3012), 2507 (HXB2 2571) nt in length respectively.

**Figure 1:**
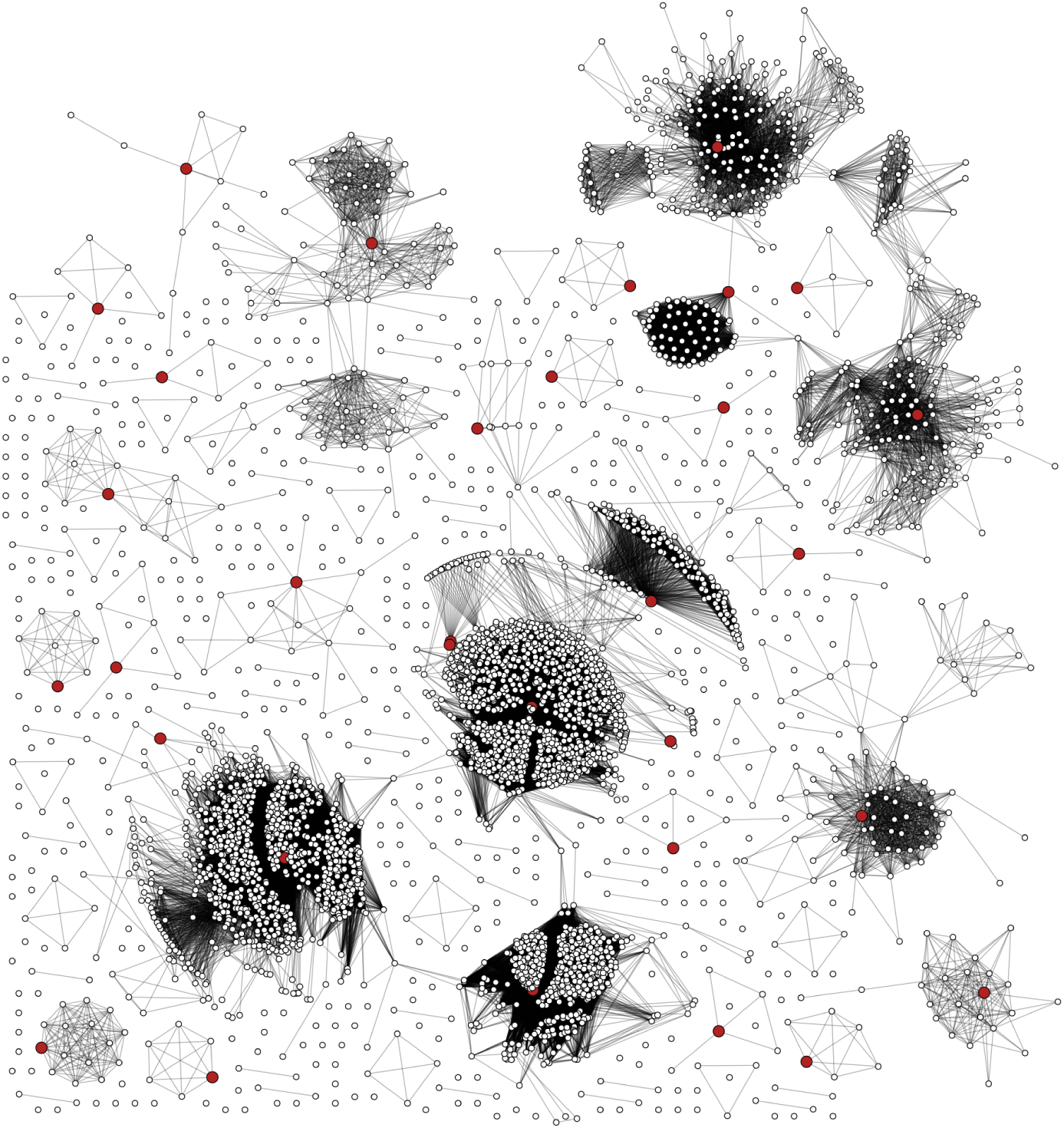
A network diagram representing the clustering of HIV-1 near full-length sequences into communities of genetically similar genomes. Each point (node) represents an HIV-1 genome sequence, and each connection (edge) indicates that the respective nodes have a pairwise *k*-mer distance below the threshold. Larger (red) nodes represent sequences with the maximum degree-size centrality in their respective network communities; these sequences were in a multiple sequence alignment to generate a draft consensus genome sequence.

### Evolutionary History of HIV-1 Group M

We employed a relaxed molecular clock analysis as implemented in the least-squares dating program LSD [19] to estimate the divergence times and substitution rate estimates across the HIV-1/M genomes. We found evidence of substantial variation in estimated root dates among windows, with an estimated mean tMRCA of 1922 (Fig. 2). Estimates were significantly autocorrelated (Durbin-Watson Test, *p* < *0.05*), which was more consistent with a genome-level effect than uncertainty in phylogenetic reconstruction and dating. We show that the most recent estimates were obtained from windows within the *env* gene region, which has frequently been targeted in studies of HIV-1 origins [2, 3].

**Figure 2:**
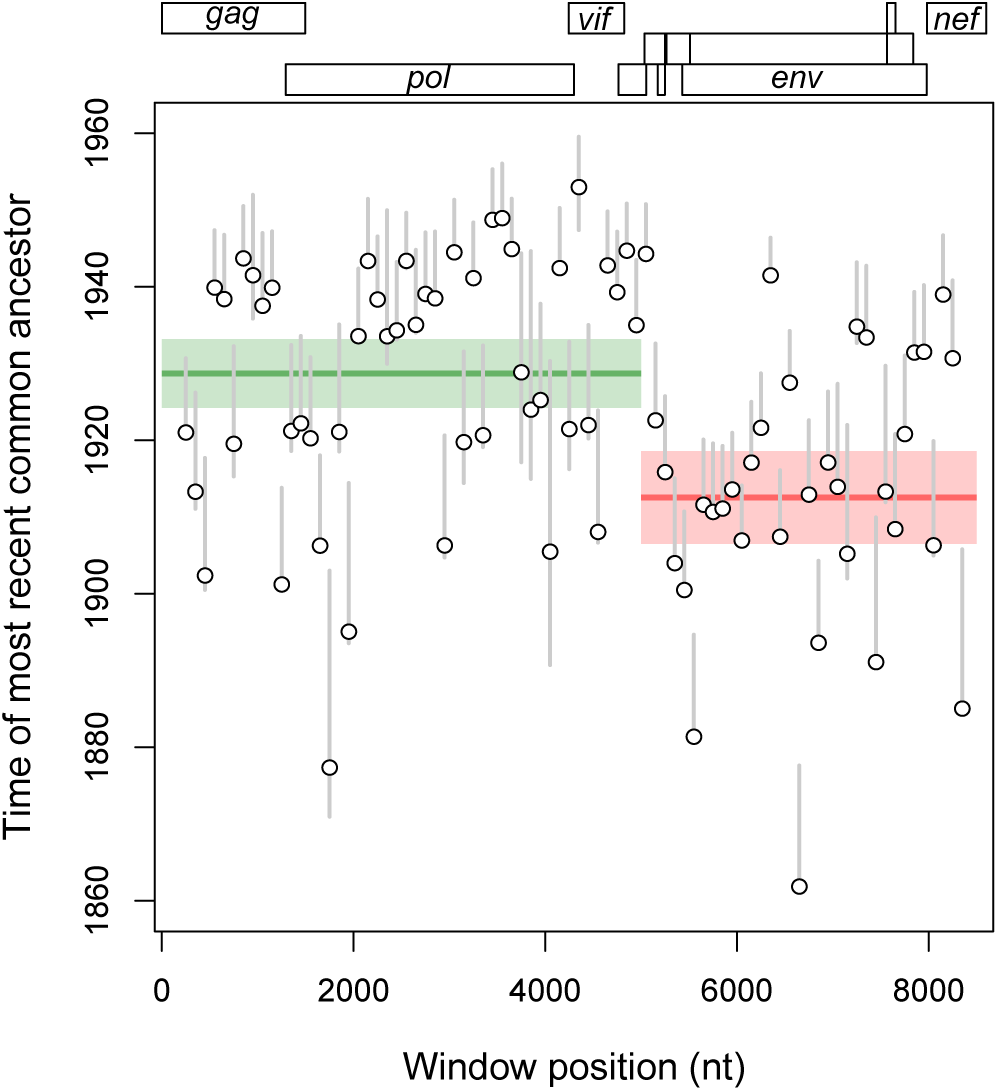
Origin of HIV-1/M based on the LSD method. The y-axis represents the estimated root dates (tMRCAs) for sequences in each window. The x-axis represents the genomic coordinates with each position corresponding to a gene region on the HIV-1 genome map at top of the boxplot. Each point represents data for a sliding window of 500 nucleotides per sequence. The grey lines represent the 95% confidence interval for each date estimate. The green and red lines indicate the breakpoint fragments computed using a piecewise linear regression model.

Our results also demonstrate an appreciable degree of variation across windows for the estimated nucleotide substitution rates (Fig. 3). The regions with the highest substitution rates generally coincide with the regions that were estimated to have the most recent dates. However, rates from the latter half of the *env* gene (about 7000 to 8000 in our coordinate system) were more similar to rate estimates obtained in *gag* or *pol*. This region (*env*) has been established as the most variable region across HIV genomes [31]. We calculated the relationship between the genomic window and estimated root dates by fitting a piecewise linear regression model to the data. Estimates using a piecewise regression model supports the existence of at least one recombination breakpoint in the HIV-1/M genome with intercepts around 1929 (95% CI: 1924 – 1933) for the younger fragment and 1913 (95% CI: 1906 – 1918) for the older fragment (Fig. 2).

**Figure 3:**
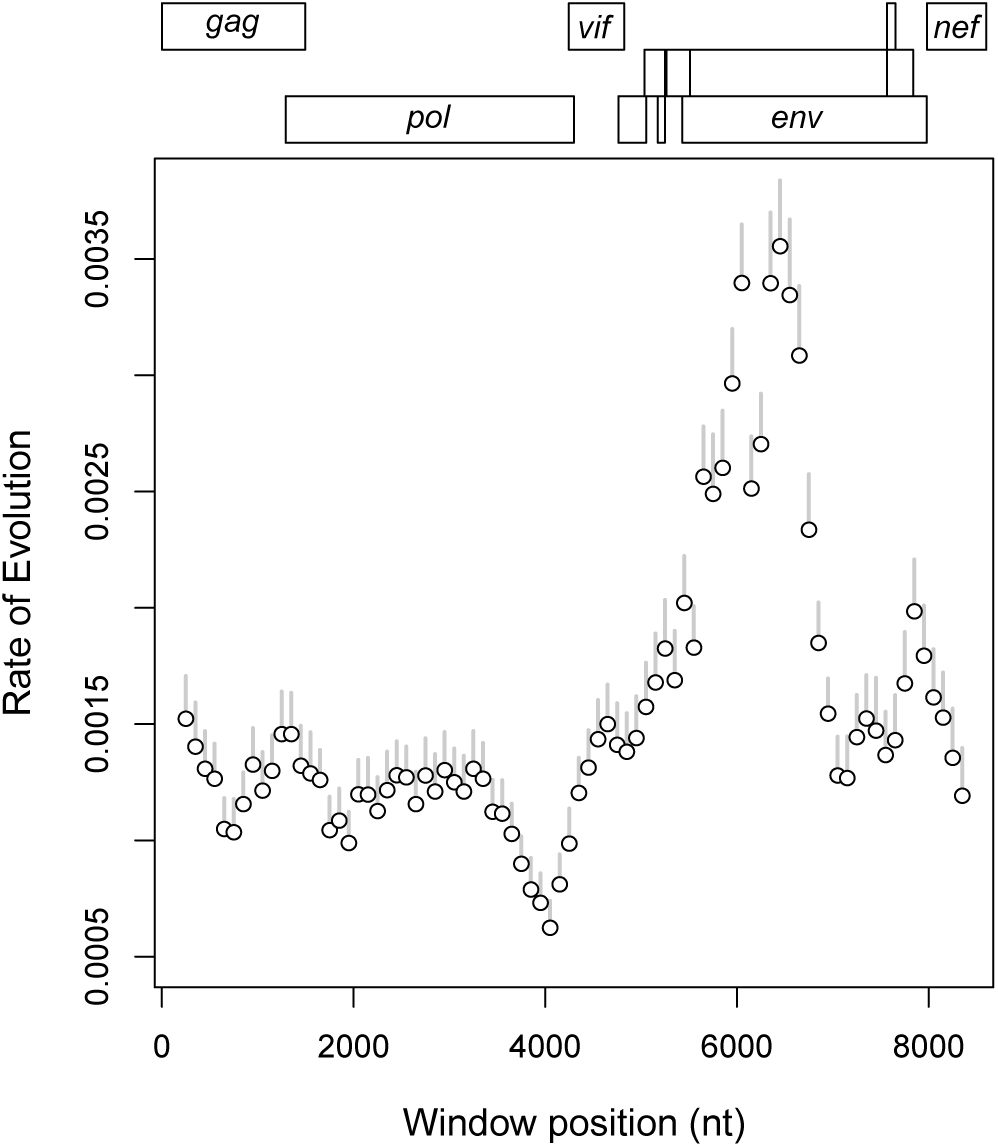
HIV-1/M substitution rate estimates based on the LSD method. The y-axis represents rates of substitution for sequences in each window. The x-axis represents the genomic coordinates with each position corresponding to a gene region on the HIV-1 genome map at top. Each point represents data for a sliding window of 500 nucleotides per sequence. The grey lines represent the 95% confidence interval for each rate estimate.

### Selection Analysis

Purifying selection occurs when lineages carrying amino acid substitutions at particular sites have reduced fitness. Thus, purifying selection can be detected by comparing site-specific non-synonymous (dN or *β*) and synonymous (dS or *α*) substitution rates, where *β* − *α* < 0 indicates purifying selection, *β* − *α* ≈ 0 indicates neutral evolution, and *β* − *α* > 0 indicates diversifying selection [32]. Purifying selection can skew dated-tip analyses because the removal of amino acid substitutions by selection causes the model to underestimate branch lengths, resulting in more recent estimates of the tMRCA. We observed substantial variation in estimates of *β* − *α* across subsets of codon sites corresponding to window alignments in the major genes of the HIV-1/M genome (*n* = 132 codons per gene, Supplementary Fig. S5). For example, the proportions of codon sites under significant (*α_p_* = 0.05) purifying selection were 0.182, 0.121 and 0.265 for *env*, *gag* and *pol*, respectively. However, we found no significant mean difference in *β* − *α* between *env* and the other two major genes (Wilcoxon test, *p* = 0.192).

## Discussion

Although HIV-1 has a high effective rate of recombination [12] and inter-subtype recombination is prevalent [13], studies reconstructing the early history of the HIV-1/M pandemic have generally ruled out the potential effect of recombination as a simplifying assumption. There is growing evidence however that recombination has played an important role in the early evolution of HIV-1/M, and recombinant genomes have been documented between recognized HIV-1 subtypes and highly divergent unique HIV-1 lineages [16, 33]. In addition, there has been growing controversy regarding the assignment of HIV-1 variants to a fixed nomenclature of pure subtypes and CRFs when our current understanding of the early and current global evolution of the virus at the genomewide level may be incomplete [15, 33–35]. Using a sliding window method on a special multiple alignment of near full-length genomes, we found evidence of substantial variation in estimated root dates (tMRCAs) among windows. Across windows, the median date estimates averaged about 1922 (range 1861 – 1948), which is consistent with previous work that places the origin of HIV-1/M within the first three decades of the 20^th^ century [2, 3, 5]. Our median tMRCA estimates were significantly autocorrelated along the genome, suggesting that the variation could not be entirely explained by uncertainty in phylogenetic reconstruction or sampling variation. Instead, this pattern implies a deterministic effect of the region of the HIV-1 genome being analysed on estimates of the tMRCA.

A significant influence of genomic regions on estimates of tMRCA can be explained by either recombination or selection, and it is possible that both processes have played some role. First, the observed pattern in tMRCA estimates may be the residual signature of at least one recombination event that occurred early in the evolutionary history of HIV-1/M, preceding the rapid expansion of this group from central Africa to the rest of the world. Put another way, the evolutionary history of HIV-1/M would be more accurately modeled by two phylogenies instead of one, where the phylogeny relating the 5’ portion of the genome has a more recent common ancestor. Second, this outcome may be the result of different levels of purifying selection in the HIV-1 genome [36]. Purifying selection reduces the observable rate of molecular evolution by removing deleterious mutations at conserved sites [37]. This greater constraint on tolerated substitutions shortens the time scale on which a molecular clock analysis is susceptible to saturation, where the number of substitutions between two sequences is so extreme that they become indistinguishable from completely unrelated sequences. Saturation of a conventional molecular clock sets an upper limit to how far back we can extrapolate in time, such that a pair of sequences that have diverged for hundreds of years are indistinguishable from their ancestors that have diverged for only decades. The HIV-1 *gag* and *pol* genes on the 5’ side of the genome tend to be more evolutionarily conserved than the *env* and *nef* genes that are encoded closer to the 3’ end [38]. Hence, the more recent tMRCA estimates from *gag* and *pol* are consistent with greater purifying selection filtering out mutations at conserved sites.

Here we enumerate the evidence from our study that support an early recombination event as a more parsimonious explanation for the distribution of tMRCA estimates along the HIV-1 genome. First, the LSD estimates of the molecular clock we obtained from *nef* and the portion of *env* encoding gp41 were comparable to estimates from *gag* and *pol* (Fig. 3). We did not observe a concomitant increase in tMRCA estimates associated with gp41 and *nef* that would signify a similar effect of purifying selection in these regions. Second, our alignment largely excluded several major targets of diversifying selection, including the hypervariable loops in *env*, because these regions are problematic for generating a multiple sequence alignment across the global diversity of HIV-1/M. Indeed our analysis of site-specific selection (Fig. S5) indicated that the majority of codon sites from the region of the filtered alignment corresponding to HIV-1 *env* were evolving under levels of purifying selection comparable to sites in *gag* and *pol*. Third, the sliding window alignments do not exhibit a clear signature of saturation. For instance, Supplementary Fig. S6 plots the numbers of transitions and transversions between pairs of aligned sequences against a genetic distance corrected for multiple hits — a conventional method for detecting saturation [39] — for an arbitrary subset of window alignments. This is consistent with recent work using a more complex time-calibrated approach, which did not find evidence of saturation in HIV-1 sequences [40].

It is important to note that we only used data that were available in the public domain, and that our data collection does not represent a random sample of the global distribution of HIV-1/M. However the primary objective was to sample the ancestors of HIV-1/M and our study does not draw any conclusions about the present global distribution of the virus. We postulate that recurrent introgression between countries of HIV-1 genomic variation through inter-subtype recombination limits the effect of sampling location, and that focusing the molecular clock analysis on relatively narrow windows of the genome instead amplifies the influence of sample collection dates. Our results support the hypothesis that the current global diversity of HIV-1/M may be a product of at least one recombination event between two or more ‘ancient’ variants, with an older fragment encoding the *env* and *nef* genes. It is tempting to speculate that an earlier origin of *env* may have been associated with the role of the envelope glycoproteins in adapting to the antibody-mediated immune response of human hosts, but this hypothesis needs to be substantiated by further investigation. Our work demonstrates a novel approach to incorporating genome-scale variation in reconstructing the evolutionary history of HIV-1, instead of focusing on a specific region of the genome. With growing availability of more cost-effective and rapid sequencing technologies and software tools for processing the resulting outputs, it is becoming more feasible to analyze the evolutionary history of HIV-1 on a genome-wide scale. In combination with appropriate phylogenetic tools, these data will inevitably advance our knowledge on the origin and evolution of HIV-1. For instance, we anticipate that the large-scale genome sequence analysis of HIV-1/M in Africa currently being conducted by the Phylogenetics and Networks for Generalized HIV Epidemics in Africa consortium (PANGEA-HIV) [20] will deliver significant new insights into the evolution of this virus.

## Acknowledgements

This work was supported in part by the Government of Canada through Genome Canada and the Ontario Genomics Institute (OGI-131) and by the Canadian Institutes of Health Research (CIHR grants PJT-155990, PJT-156178, FRN-130609, BOP-149562).

## Supplementary Figures

**Figure S1:**
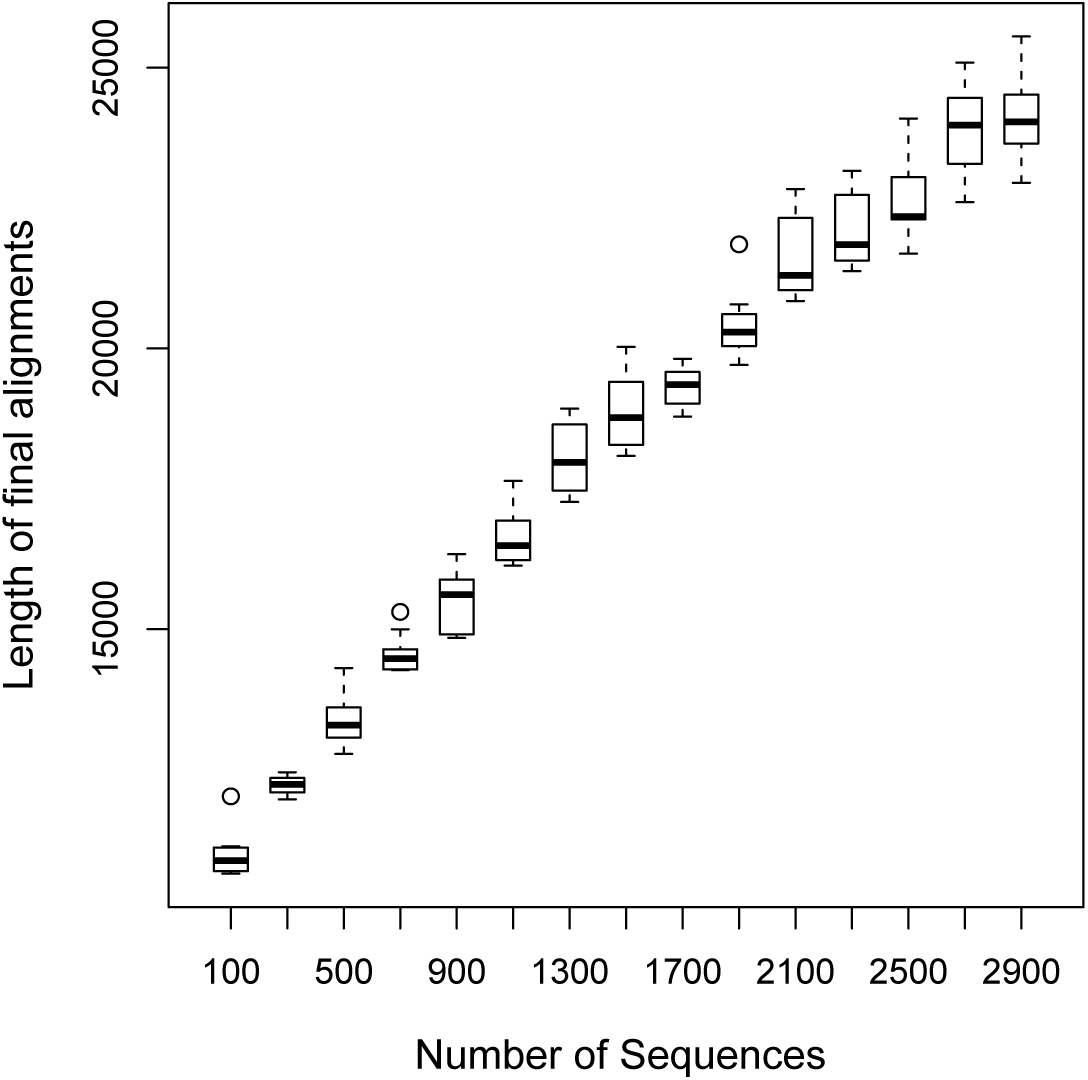
Boxplot showing the relationship between the number of HIV-1/M sequences and the length of the final alignment. The alignments were generated using the MAFFT program. The y-axis represents the the length of the final alignment (nt) generated, while the x-axis represents the number of sequences used in each run.

**Figure S2:**
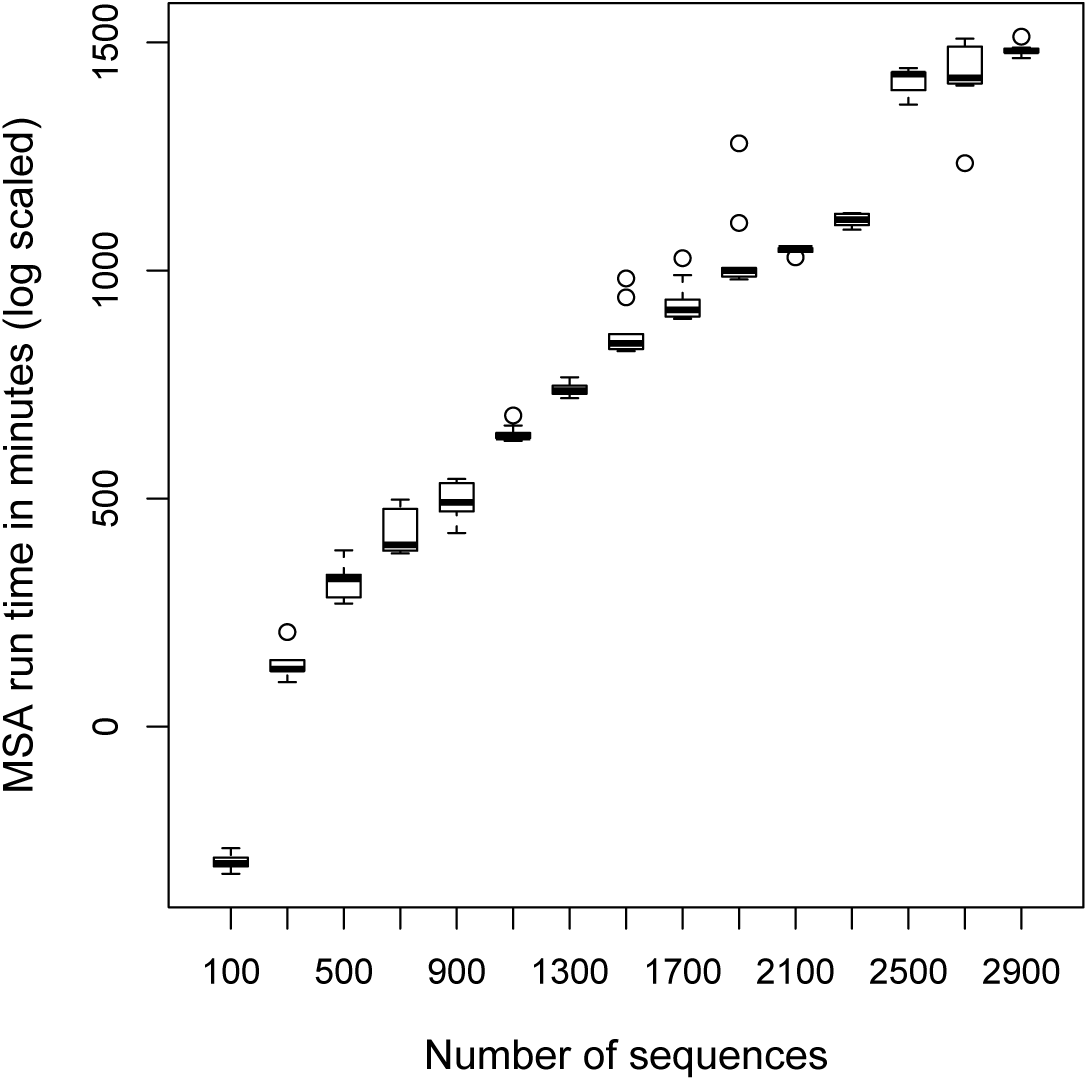
Boxplot showing the relationship between the number of HIV-1/M sequences and the time it takes to generate an alignment. The alignments were generated using a conventional multiple sequence alignment program called MAFFT. The y-axis represents the run time in minutes (log scaled) and the x-axis represents the number of sequences used in each run.

**Figure S3:**
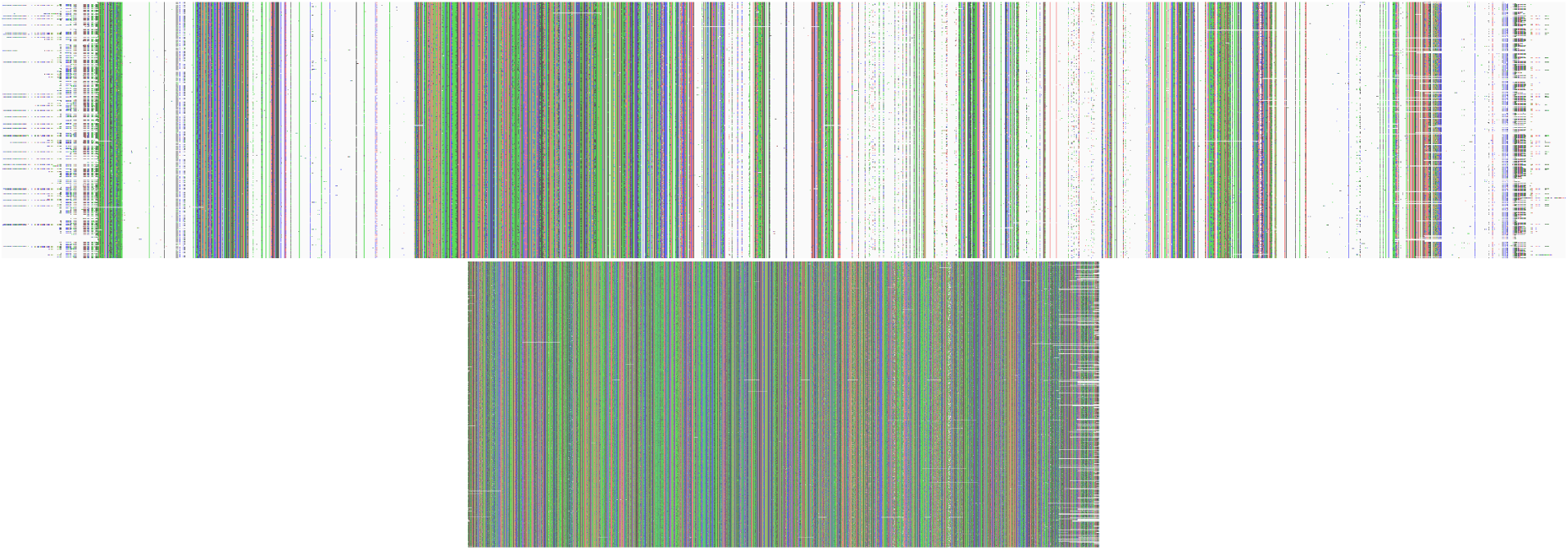
Snapshots of unprocessed and processed HIV-1 group M sequence alignments. These image files were generated using the program AliView (http://ormbunkar.se/aliview). (top) An alignment of 3,500 HIV-1 genome sequences produced by MAFFT. Note that some individual nucleotides are not visible at this resolution. (bottom) An alignment of all 3,900 genome sequences after pairwise alignment processing.

**Figure S4:**
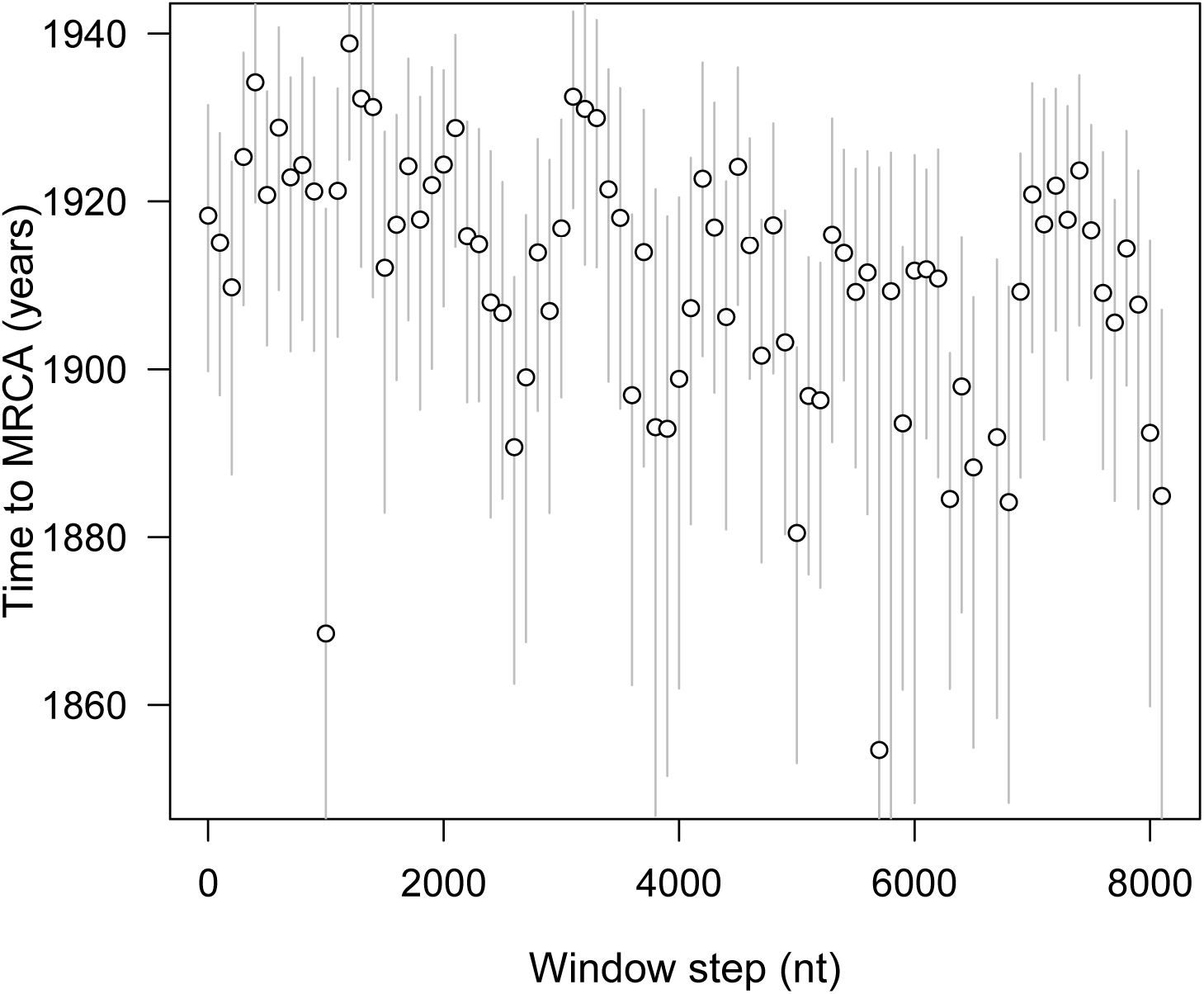
Origin of HIV-1 group M based on the Bayesian method. The y-axis represents the estimated root dates (tMRCAs) for sequences in each window. The x-axis represents the genomic coordinates with each position corresponding to a gene region on the HIV-1 genome map at top of the boxplot. Each point represents data for a sliding window of 500 nucleotides per sequence. The grey lines represent the 95% confidence interval for each date estimate.

**Figure S5:**
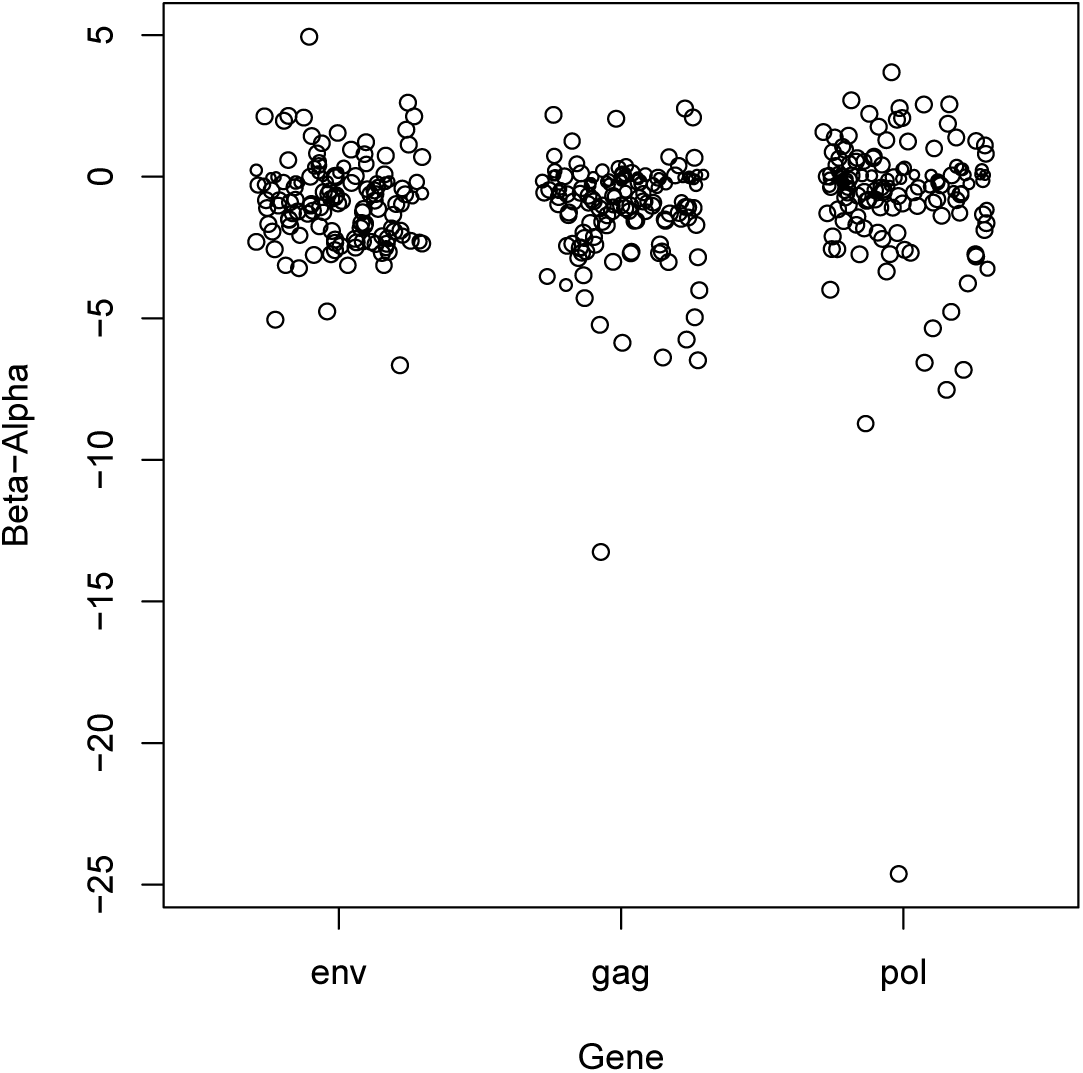
Selection analysis of the HIV-1/M major genes. The y-axis represents the mean posterior of the non-synonymous (dN) rate minus the synonymous (dS) rates per site. The x-axis represents each of the major gene

**Figure S6:**
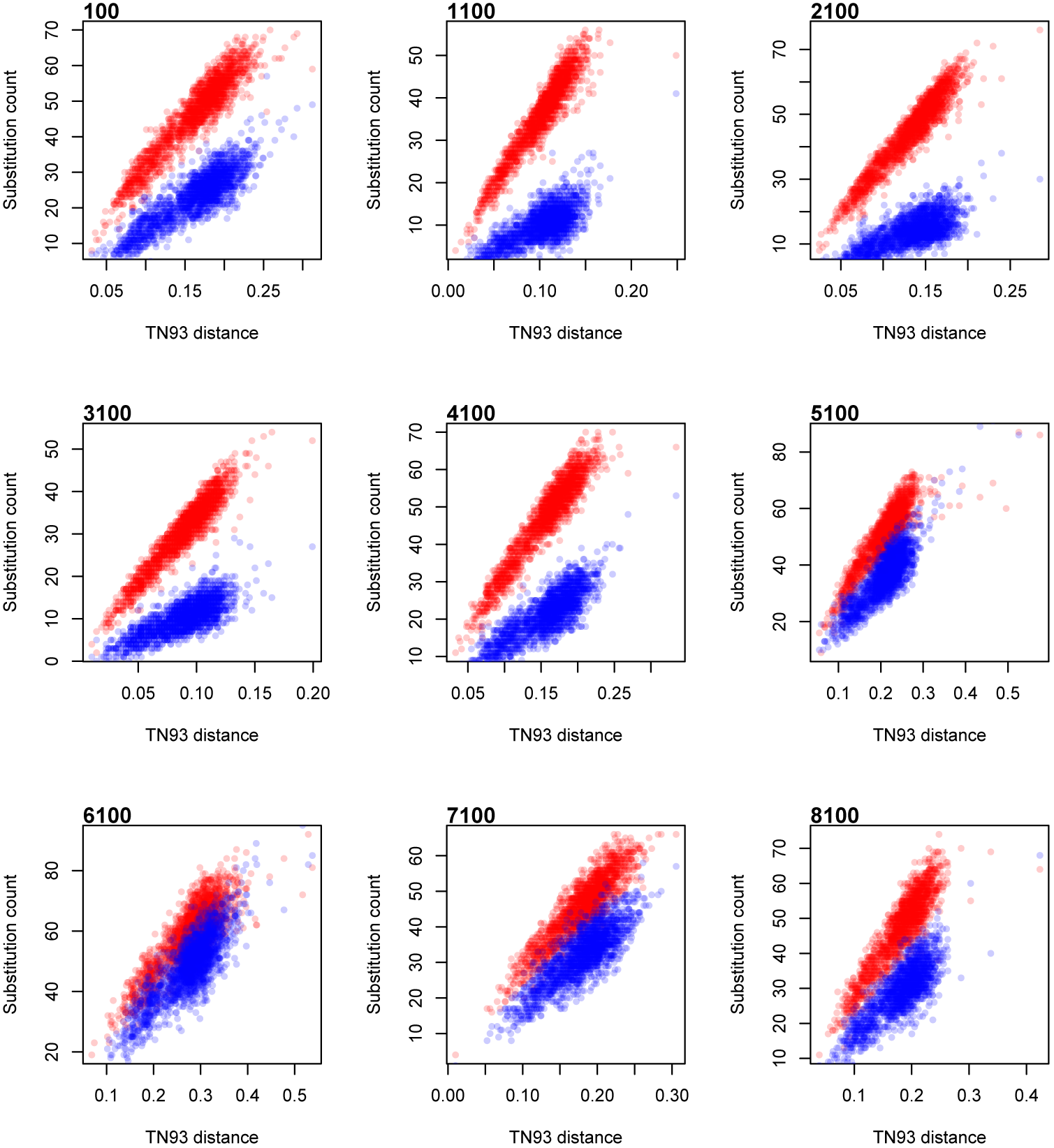
Plots of transition and transversion substitution counts against the Tamura-Nei (TN93) genetic distance in nine window alignments. The upper label on each plot denotes the leftmost position of each window alignment relative to the consensus genome sequence. For each window alignment, we generated a random permutation of the sequences and then for each pair of sequences, we counted the number of transitions (red) and transversions (blue) and the TN93 distance.

